# Fast or slow - light climate modulates intra-population sinking velocities in small phytoplankton

**DOI:** 10.1101/2025.06.13.659648

**Authors:** Maximilian Berthold, Douglas A. Campbell, Rahel Vortmeyer-Kley

## Abstract

The global carbon cycle depends heavily on the carbon sequestration rates of aquatic ecosystems. Sinking of phytoplankton is a rapid mediator of carbon sequestration, because phytoplankton are globally abundant photoautotrophs that grow rapidly. Pico- and nano-phytoplankton sinking velocities vary depending on their growth state, viability, clumping, and distribution in the water column. We introduced high throughput fluorescence microscopy of well-plates, to measure sinking velocities of three diatom strains, and three cyanobacteria strains, with cell radii spanning an order of magnitude, all grown under three different light levels. Cultures were measured for sinking velocities repeatedly across their growth trajectories. Tracking multiple fluorescence wavebands allowed us to simultaneously determine sinking velocities for living vs. dead cells. Sinking velocities varied strongly across growth light levels, and across growth stages. These monoclonal cultures furthermore show distinct sub-populations of slow- and fast-sinking cells. Our results departed widely from simple Stokes’ Law estimates of sinking based upon radii and mass density of cells. Complex, heterogeneous phytoplankton communities likely show more complicated sinking patterns than are currently expressed in biogeochemical ocean models. Our well-plate microscopy approach using parallel imaging of many samples generates high-throughput measures of cell sinking at population- or community-scales, to in turn improve modelling of carbon export to deeper layers.

## Introduction

Sinking is one of the four main processes that govern phytoplankton distribution and abundance in the water column, along with growth, grazing and senescence, including viral lysis. Sinking influences the local abundances of phytoplankton cells within the water column, which in turn impacts underwater light climate and nutrient availability. Sinking phytoplankton also constitutes a large share of organic matter exported into the aphotic zone, where it is eventually sequestered as part of marine snow, which then sinks rapidly to the deep parts of the ocean [1, 2].

On a global scale, this biological carbon pump affects the amount of carbon that sinks from the euphotic zone to the deeper parts of the coastal and open oceans. The biological carbon pump is increasingly impacted by global climate change, which alters water temperatures, stratification, and mixed layer depth, as well as changing wind fields and solar insolation [2–5]. With increasing temperature smaller phytoplankton may increase in abundance and in total population biovolume [6]. Picocyanobacteria are thus seen as future sentinels of a warming ocean, but other phyla, like diatoms, may also experience community shifts towards smaller cells [6, 7]. Prospective changes in abundances of different phytoplankton size classes will affect the total amount of fixed and exported carbon in the upper part of the ocean, impacting the food web, microbial loop and biological carbon pump. Thus, it is critical to measure, and understand how nano-to picophytoplankton communities sink through the euphotic and aphotic zone of the upper water column, to improve global biogeochemical ocean models and ultimately better understand global carbon export [2, 8, 9].

Sinking velocity is actively regulated, at least in some larger cells [10], and through accumulation of gas vesicles in some cyanobacteria [11]. Furthermore light availability and concomitant carbohydrate production and consumption [10, 12] dynamically influence cell density, although such density changes may only significantly influence sinking velocities in cells with larger radii [10].

Phytoplankton sinking velocities also change over the course of the life cycle, and through long term adaptations of cell composition or shape [1, 10, 13–15]. Previous studies of phytoplankton sinking velocities, in the absence of mixing, have taken into account phytoplankton shape and inner structure [16, 17], cell size and growth conditions [18–20], temperature [21] and the density [22] of the surrounding fluid.

Positive buoyancy regulation, and interactions of light with sinking velocities have also been considered for freshwater ecosystems ( [23] and references cited therein). Finally, phytoplankton sinking is disturbed by external hydrodynamical forces in the ocean including upwelling, downwelling, turbulent mixing processes and stratification, which are themselves under long-term climate influences (e.g. [24–28]).

Despite this empirical knowledge on phytoplankton sinking, for modelling purposes phytoplankton sinking velocity has been approximated using only cell radii and generic cellular density as inputs into Stokes’ law, or modifications thereof [17, 22, 25, 26, 28]. Stokes’ law is, however, a purely physical simplification that does not take into account the importance of biological mechanisms in particle sinking [29]. Furthermore, mass density data for living phytoplankton, required for Stokes’ estimates, are often missing or are inferred from simplified biovolume conversions. Even such basic cell size effects are neglected in 13 of 19 climate models that include a biogeochemical component, making realistic predictions of future carbon export difficult [9].

Instead, for modelling a fixed sinking rate is often assumed for functional groups of plankton or types of organic material (e.g., detritus, fecal pellets, or dissolved organic material), when indeed sinking of living phytoplankton or dead organic material is even considered in biogeochemical ocean models (e.g. [30–32]).

This limited incorporation of phytoplankton sinking into biogeochemical ocean models makes it important to efficiently quantify nano-to pico-phytoplankton sinking, since these functional groups have potentially increasing roles in the warming ocean. The purpose of this investigation is thus to establish and demonstrate a high-throughput approach for measuring sinking velocities for nano-to pico-phytoplankton, applicable to heterogeneous populations, and to compare our results against Stokes’ law approximations of sinking. We thereby aim to support more realistic modelling of sinking in biogeochemical ocean models. Sinking can potentially be regulated in living cells, but not in dead cells [33, 34], so our approach supports resolution of both phytoplankton size, and viability, classes, reflecting different life stages and sizes. We firstly introduce our well-plate fluorescence microscopy approach (Fig. 1A) to analyze phytoplankton sinking. We then present and discuss the results of the experimental setup with a focus on research questions:

**Fig 1.**
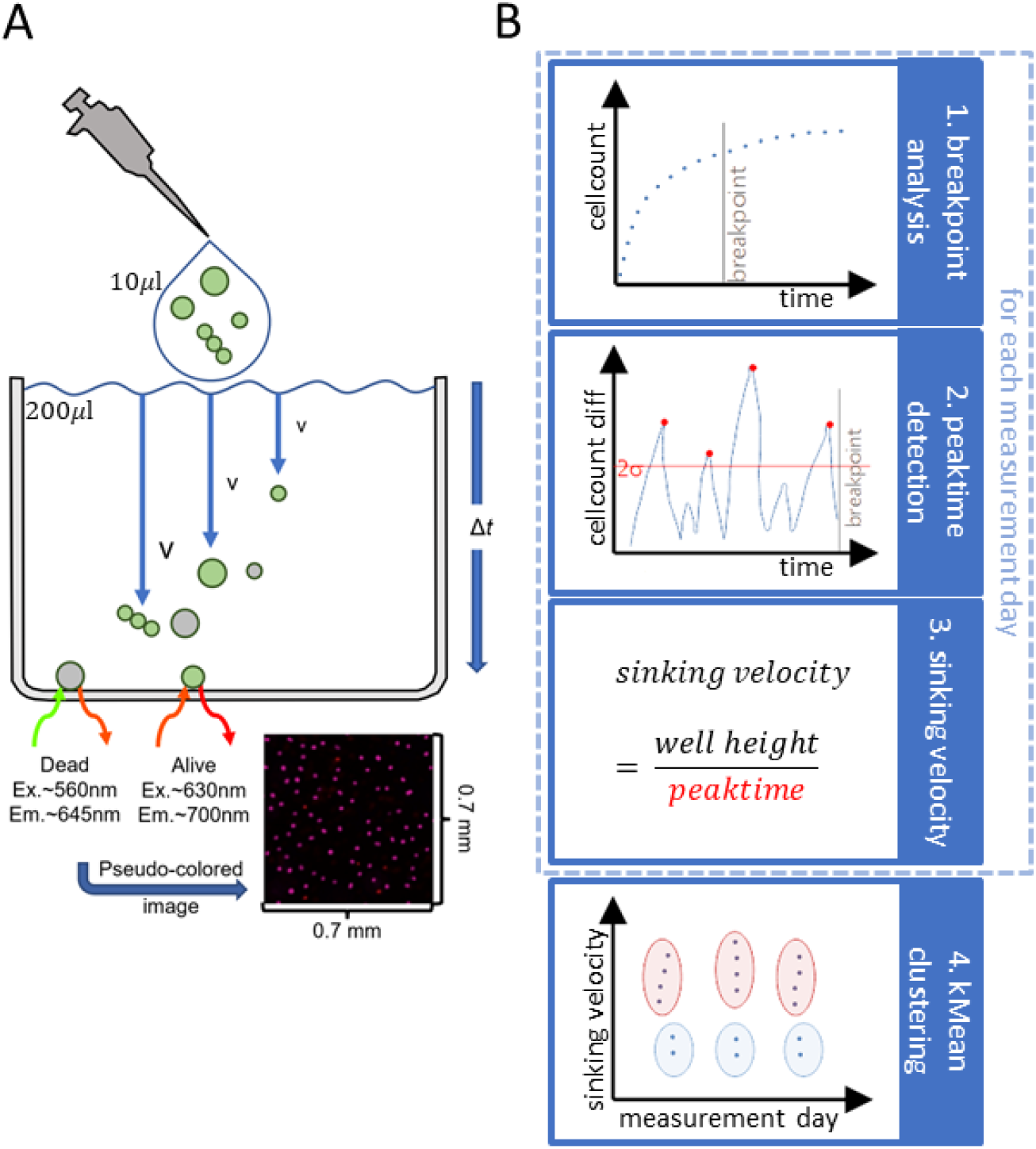
(A): Detection of cell sinking velocities in a monoclonal phytoplankton sample, but with heterogeneous cell sizes, viability states, and cell colony formations (schematic sketch of one well out of 96 wells). In the beginning of the measurement the sample is pipetted onto the surface of the well filled with respective strains media. Cells were counted at 10-min intervals as they reached the bottom of the well, with sinking velocities estimated from the rates of accumulation at the bottom (analysis of sinking velocity in Fig.1B). The dual-channel analyses allowed us to distinguish living diatoms or cyanobacteria (green symbols; pigment fluorescence; no propidium iodide fluorescence); from dead or dying diatoms or cyanobacteria (gray symbols; no or residual pigment fluorescence along with propidium iodide fluorescence). The image was automatically pseudo-colored by the ImageXPress Software (Molecular Devices), red for cells stained by propidium iodide, and pink for pigment fluorescence. The example image was taken from a time course of *Synechocystis* sp. sinking in a single well of a microwell plate at one time point. (B): Steps of raw data analysis segregated by cell viability and by well: 1. breakpoint analysis [37]: separation of sinking and plateau phase; 2. peaktime detection: calculating time-lagged differences in cell counts between neighbouring time points of the sinking phase; detection of significant peaks in this signal; significant peaks higher than twice the standard deviation of the average time-lagged differences of the cell count; 3. sinking velocity calculation: different sinking velocity per well represent different parts of a population sinking out; 4. kMean clustering: separation into fast- and slow-sinking sub-populations;

1. How does sinking velocity of phytoplankton vary depending on their stage of life and their size?
2. Does light level during phytoplankton growth impact sinking velocities?
3. Do measured sinking velocities of phytoplankton depart from Stokes’ law estimates?

An unexpected challenge in this study was a need to deconvolute multiple classes of cellular sinking velocities (Fig. 1B), because even within single-strain cultures we resolved sub-populations showing different sinking velocities, which varied over growth trajectories.

Throughout the text, references to supplemental material are marked with S.

## Material and Methods

### Cell properties and growth conditions

As picocyanobacteria account for approximately 25% of global aquatic primary production, while diatoms account for 40% [35, 36], this study therefore included picocyanobacteria (*Synechocystis* sp. - PCC6803; *Cyanobium* sp. - CZS25K; *Synechococcus* sp. - CCMP836) and diatoms (*Thalassiosira weissflogii* - CCMP1336; *Thalassiosira pseudonana* - CCMP1335; *Minidiscus variabilis* - CCMP495) (for details cf. Table 1). The strains spanned diameters from 1.3 to 20.0*µ*m, thus falling within the pico-to nano-phytoplankton size classes.

**Table 1.**
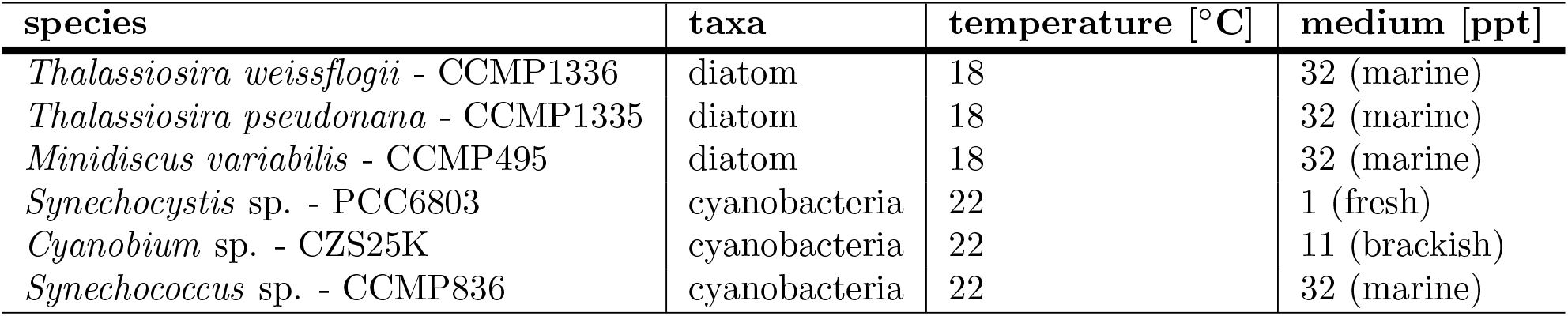
Diatoms and cyanobacteria studied and their growth conditions.

Diatoms were grown in full marine f/2 media at 18°C, whereas cyanobacterial strains were grown in BG11 media with varying salinities depending on on the strain isolation location (*Synechocystis* sp. freshwater, *Cyanobium* sp. brackish, *Synechococcus* sp. full marine) at 22°C (cf. Table 1). Pre-cultures were grown under a 12:12 Light:Dark cycle, at a light level of 60 to 80 *µ*mol photons m^−2^ s^−1^ (abbreviated as *µ*E on figures) on a shaker, and transferred, every seven days into fresh media.

For the experimental culture growth, wells of 12-well plates were loaded with 4ml of respective culturing media, along with a sterilized magnetizable microstir bar. Wells were then inoculated with 1ml of cyanobacteria or diatom culture. The well plates were then placed at growth temperatures of 18°C for diatoms and 22°C for cyanobacteria, under a 12:12 Light:Dark cycle, at growth light levels of 300, 100, or 30*µ*E. These light levels were chosen as representative of the upper to lower euphotic zone. Four replicate well cultures were grown for each combination of strain and growth light. Strains remained monoclonal within their respective wells and were resuspended once to twice a day using the magnetic microstir bars at less than 100rpm.

### Sinking Velocity analysis

To estimate sinking velocities over culture growth trajectories, we took samples from the well-plate cultures daily for a week from well cultures of the monoclonal strains listed in Table 1, and loaded them in a 96-well plate.

For determinations of cell sinking velocities we used a plate-based cytometer equipped with an inverse fluorescence microscope (Molecular Devices Pico image analyser) and its onboard ImageXpress software (Molecular Devices, San Jose, CA, USA) to analyze the fluorescent images. The instrument allows Z axis focussing through the well depth, which we used to capture images of cells on the well bottom, to generate total cell counts, average and total cell fluorescence, as well as average and total cell area on the bottom of the well.

The image capture area was in the center of each well (0.48mm^2^ of 34mm^2^), to limit wall effects interfering with sinking. Cells were differentially imaged by two fluorescence channels. Living cells were detected from their auto-fluorescence using the ‘Cy5’channel (Excitation 610-650nm, Emission 675-720 nm). 50*µ*l culture were stained with 20*µ*l propidium iodide (15*µ*M) to sample only dead and dying cells [38], detected through fluorescence using a ‘TxRed’ channel (Excitation 535-585nm, Emission 610-680nm).

Subsequently a 10*µ*l aliquot was pipetted onto the surface of a 96-well plate filled with 200 *µ*l of respective strain media right before the start of the experiment. Cells were scored as ‘dead’ if only fluorescence from propidium iodide was detected, while cells were scored as ‘dying’ if both auto-fluorescence and propidium iodide fluorescence was detected from the cell. The data sets for the dead and the dying cells were small and thus noisy, so we pooled the dead and dying cell data into a ‘dead and dying’ category to improve analyses. It is important to note that cells were not fixed with chemicals, because living and dead cells might show distinct sinking properties. We were thus able to track two distinct life stages within each culture.

We then took fluorescence microscopy images of the well plate repeatedly every 10min, over a time course of 10h. The onboard ImageXpress software analysed total cell counts, average fluorescence per cell, total cell fluorescence, as well as average cell area and total cell area on the bottom of the well, based on pixels falling within pre-defined size ranges for respective strains. We then analysed the data using R with RStudio [39]. A graphical overview of the workflow of the analysis can be found in Fig.1B.

For each well, on each day, we segregated data into counts of living and dead and dying cells. Within these sub-populations the cell count data typically showed an initial phase of rapid increase of cell numbers reaching the bottom over time, identified as sinking phase, followed by a subsequent plateau phase (cf. Fig. 1B Step 1). We applied breakpoint analysis for living, dead and dying cells for data from each day, from each well, to separate data into sinking vs. plateau phases using using the breakpoints-function (R package strucchange, [37]) (cf. Fig. 1B Step 1). For the sinking phase we then calculated time-lagged differences of cell counts between neighbouring time points to detect peaks of cell sub-populations falling simultaneously, either due to similar size, or life stage. We selected only peaks that were twice the standard deviation of the average cell difference between sequential measurement times for the well (cf. Fig. 1B Step 2). A potential problem for the peak detection was the loss of signal for propidium iodide-stained cells over time, which would result in loss of detection of dead and dying cells. We addressed this by only performing peak detection up to the time at which the maximum count of dead cells per well bottom was achieved, usually within three hours of the time course. For dying diatom cells this loss of signal and the possible misidentifications of dying cells as either living or dead ones, leads to larger errors in case of lower cell numbers. In particular the data set for cells of the largest diatom size class was sparse due to lower cell numbers per field of view.

We used the time of the occurrence of a peak to calculate the sinking velocity of the respective cells based on well height and time since the start of the sinking experiment. These peak detections resulted in several sinking velocities per well representing different sub-populations (Fig.1B Step 3). During the analyses we noted that cells fell into fast (within the first hour) or slower sinking cell sub-populations (within the first 2-3 hours). We used a cluster-analyses (kMeans) to create this additional grouping variable for modelling purposes (Fig.1B Step 4). A generalized additive model (GAM) was then generated for functional groups of cyanobacteria, diatoms strains with diameter less than 10*µ*m and diatoms strains with diameter larger than 10*µ*m based on growth light, viability state (living vs. dead and dying) and slow vs. fast sinking velocity clustering groups.

The code used to analyse the data is public available at https://github.com/maxberthold/PhytoplanktonSinkVelocities.

### Sinking according to Stokes’ law

Sinking velocities of spherical objects falling under the case of Reynolds numbers smaller than 1 can be described by Stokes’ law. Several studies have used Stokes law or a modified version of Stokes’ law to estimate sinking velocities of plankton and marine particles [17, 22, 34].

For the spherical cyanobacteria we calculated the sinking velocity using Stokes’ law according to [17]

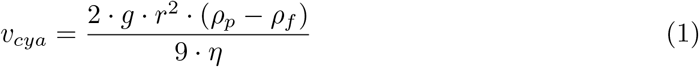

with *g* = 9.81m *·* s^−2^ as gravitational acceleration, *ρ*_*p*_ as density of the sinking particle and *ρ*_*f*_ as density of the surrounding fluid (*ρ*_*f*_ = 1010 kg *·* m^−3^ according to [44]). We assume that the cyanobacteria have the same density as the cytoplasm (*ρ*_*p*_ = 1065 kg *·* m^−3^ according to [17]). *r* is the radius of a cyanobacteria. The dynamic viscosity *η* for water is given according to [45] as temperature *T* depending parameter as 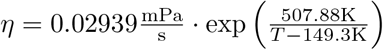 with *T* given in K (We assume here 20°C).

For the diatoms we used a combination of the modified version of Stokes’ law give in [17] and the form resistance Φ given in [22] to approximate the theoretical possible sinking velocities *v*_*dia*_:

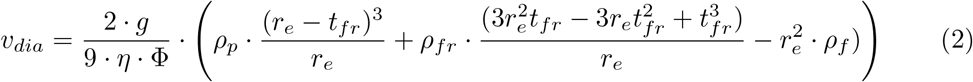

*r*_*e*_ is the effective radius of a sphere of the same volume as the cell volume *V* (*r*_*e*_ = (3 *· V/*4 *· π*)^1*/*3^). *t*_*fr*_ is the thickness of the frustule with the density *ρ*_*fr*_ = 1800 kg *·* m^−3^ according to [17]. The parameter Φ is the form resistance that describes the difference of an object from a sphere. A sphere has Φ = 1. According to [22] the cylindrical plankton species can be approximated as ellipsoid with semi-axis *a, b* and *c*. With this Φ yields 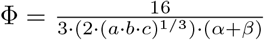. The parameter *α* and *β* describe if a particle sinks along its horizontal axis or its vertical axis.

For horizontal sinking *α* and *β* are given as 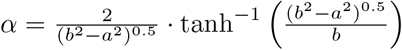 and 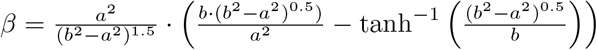 with *a* = *c* = *r* and *b* = *h/*2 and *r* is the radius and *h* the height of the cylindrical diatom.

For vertical sinking they are chosen as 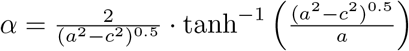 and 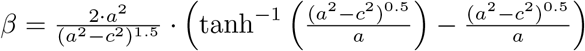 with *a* = *h/*2 and *b* = *c* = *r*.

Depending on the orientation of the particle the Stokes’ predicted sinking velocity differs, therefore we compared our experimental measurements for cylindrical plankton, where we could not estimate the sinking axis, to a mean of the horizontally oriented sinking velocities and vertically oriented sinking velocities.

Cell properties of the species studied as well as sinking velocities according to Stokes’ law (Equation 1 and 2) are given in table 2.

**Table 2.** Properties and Stokes’ predicted sinking velocity of diatoms and cyanobacteria studied. Shape and corresponding dimension data according to [40–42] (*Thalassiosira pseudonana* and *Thalassiosira weissflogii* ), [17, 43] (*Minidiscus variabilis*) and own measurements (*Synechocystis* sp., *Cyanobium* sp. and *Synechococcus* sp.). The sinking velocity is calculated according to Eq. 1 and 2. For diatoms the velocity range includes vertical to horizontal sinking velocities.

| species | shape | diameter [$\mu\text{m}$] | height [$\mu\text{m}$] | frustule thickness [$\mu\text{m}$] | Stokes' sinking predicted velocity [$\text{mm}\cdot\text{min}^{-1}$] |
| --- | --- | --- | --- | --- | --- |
| *Thalassiosira weissflogii* | cylinder | 13.0 | 10.0 | 0.75 | 1.581-1.665 |
|  | cylinder | 16.0 | 14.0 | 0.75 | 2.236-2.297 |
|  | cylinder | 20.0 | 15.0 | 0.75 | 2.731-2.891 |
| *Thalassiosira pseudonana* | cylinder | 5.0 | 3.3 | 0.1 | 0.105-0.144 |
|  | cylinder | 7.0 | 4.6 | 0.1 | 0.171-0.185 |
| *Minidiscus variabilis* | sphere | 3.9 | - | 0.05 | 0.055 |
| *Synechocystis* sp. | sphere | 2.3 | - | - | 0.009 |
| *Cyanobium* sp. | sphere | 1.5 | - | - | 0.004 |
| *Synechococcus* sp. | sphere | 1.3 | - | - | 0.003 |

## Results

### Impact of growth stage and light on sinking

We measured the sinking velocities of the diatoms and cyanobacteria in Table 1 following the steps in Fig. 1B. The analyses of the measured sinking velocities for both living and dead and dying cells show two distinct clusters with either fast (within the first hour) or slow sinking cells (within the first 2-3 hours). Therefore, we applied kMeans function from R as cluster-analyses using three cluster to the sinking velocity data to create an additional grouping variable of fast-sinking vs. slow-sinking sub-populations (cf. Fig. 1B Step 4), representing different elements of the monoclonal populations sinking out at different peak times.

Our measured sinking velocities - with values between 0.1 to 1mm*·*min^−1^ for cyanobacteria and 0.1 to 1.8mm*·*min^−1^ for diatoms varying depending on size, viability state and light conditions (cf. Fig.2) - are in the same order of magnitude as reported in other studies e.g. [13, 14, 17, 24] and references in Table 2 in [46].

As outlined in Fig. 1A, we expected an increase in sinking velocity with increasing cell size, or with the formation of cell colonies. The majority of cells across all strains reached the bottom of the well after 120 to 180 min, confirmed by a breakpoint analysis. Over that time span the average areal fluorescence signal per cell, indicative of cell size or chloroplast size, became smaller as time progressed in some, but not all, cultures, particularly for dead cells (cf. Fig.S1 and S2). In other cases, particularly for living cells, cell sizes reaching the bottom of the well were steady or even increased (cf. Fig.S1 A-C). Additionally, the cell size dynamics during measurement changed with growth day and as expected maximal reached cell size per strain depend on growth light level (cf. Fig.S1 and S2). Thus, cell size within populations was not a strong predictor of sinking rates.

The fast-sinking and slower-sinking sub-populations show up as a gap between the two boxplots for each viability state, cell size/taxa grouping and growth light in Fig. 2, across all cyanobacterial and diatom strains with diameters *<* 10*µ*m.

**Fig 2.**
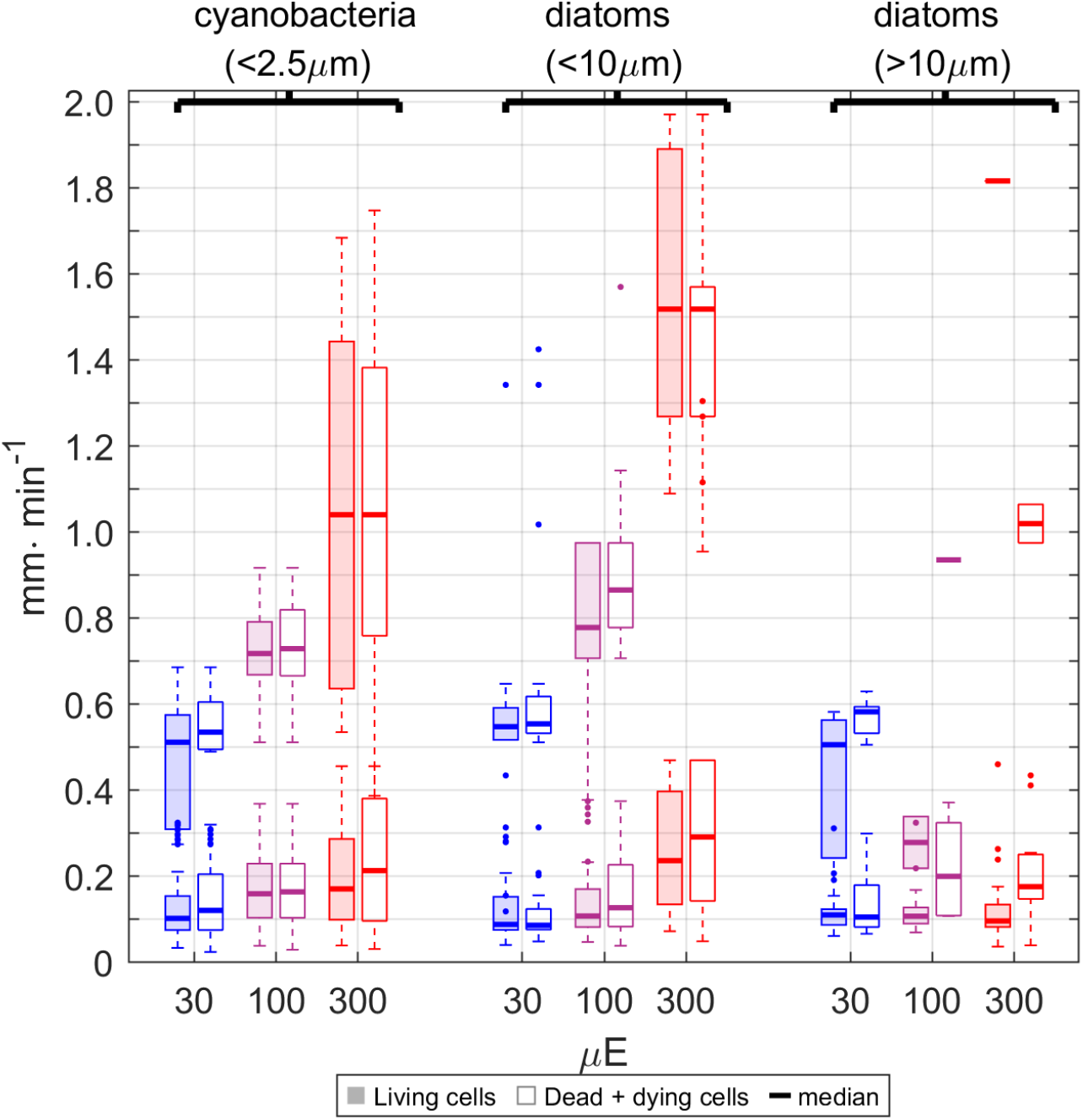
Whisker-box plot distribution of measured sinking velocities (mm· min^−1^) of cyanobacterial strains (*Synechococcus* sp., *Cyanobium* sp., *Synechocystis* sp.), small diatoms *<* 10*µ*m (*Minidiscus variabilis, Thalassiosira pseudonana*) and a diatom *>* 10*µ*m (*Thalassiosira weissflogii* ) across growth lights of 30 - 300*µ*E. Boxes are separated by viability state (shaded - living cells, empty - sum of dead and dying cells) and growth light into respective kMean-clusters. kMean-clustering consistently identified two sinking velocity clusters per light condition, size, viability and taxa groupings. The thick line indicates the median, box length 25 and 75% percentile, whiskers the 10 and 90% percentile, and single points outliers. Conditions with insufficient data show only one median value. The numbers in the brackets indicate the cell diameter.

The fast-sinking sub-populations shifted towards yet faster sinking velocities with increasing growth light Fig. 2, so the gaps between fast- and slower-sinking sub-populations increased with increasing light. At a given growth light level the diatoms with diameter*<* 10*µ*m tended to sink faster than diatoms with diameter*>* 10*µ*m, at least at growth lights at or above 100*µ*E. Interestingly, living and dead and dying cells generally showed similar sinking velocities under a given growth light, in both the faster- and slower-sinking sub-populations.

In contrast the largest diatom *Thalassiosira weissflogii* showed a decrease in sinking velocities, at least in the faster-sinking sub-populations, with increasing growth light, although there were too few observations to determine two significantly separated sinking clusters in *Thalassiosira weissflogii* under some of the growth light and viability combinations. Thus, we cannot fully confirm if there were truly two sinking clusters.

Surprisingly the largest diatom strain *Thalassiosira weissflogii* included cell populations that sank much slower than the other smaller diatom strains, and to some extent even the cyanobacterial strains, suggesting greater ability of sub-populations of this strain to actively regulate sinking velocities [47]. Active regulation of sinking of *Thalassiosira weissflogii* is further suggested because the dead and dying cells of *Thalassiosira weissflogii*, which could not regulate their mass density, generally sank faster compared to living cells. Again, cell size is not a strong predictor of measured sinking rates across the taxa range.

Following the measured sinking velocity of the fast-sinking and slower-sinking sub-populations along their growth trajectory (Fig.3) showed an increase of sinking velocity for the fast-sinking sub-populations for the cyanobacteria in the first growth days which is most prominent for high growth light (300*µ*E) (cf. Fig.3 left column). In contrast, the fast-sinking sub-population of the diatoms with diameter *<* 10*µ*m showed a decrease of sinking velocity within the first growth days which is most prominent for low growth light (30*µ*E) (cf. Fig.3 middle column). The slower-sinking sub-population of all studied species had more or less a constant sinking velocity along the growth trajectory. For the largest diatom strain *Thalassiosira weissflogii* no clear dynamics of the sinking velocity along the growth trajectory could be identified because of too sparse data for a GAM analysis especially in case of dead and dying cells (cf. Fig.3 right column).

**Fig 3.**
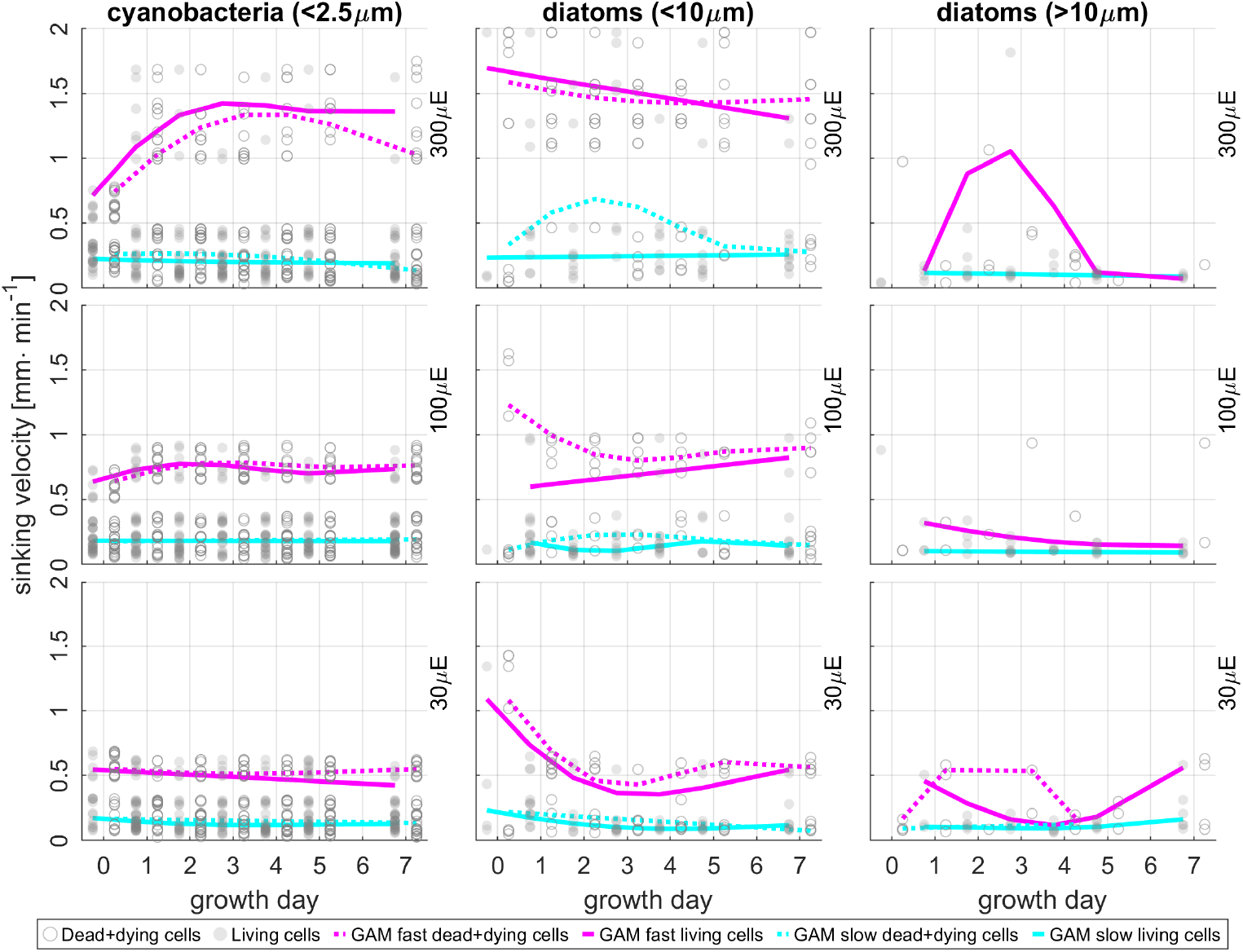
Dynamics of the measured sinking velocities of cyanobacteria and diatom strains (columns) along their growth trajectory from day 0 to day 7 (x-axis) for different cell types and viabilities (filled circles: living cells and unfilled circles: dead and dying cells; measurement dots are slightly shifted horizontally for visibility but were all measured at the same growth day.) and different growth lights (rows). The dynamics of slow-sinking sub-populations (cyan) and fast-sinking sub-populations (magenta), pooled from all cyanobacteria, diatom strains with diameter *<* 10*µ*m or diatoms with diameter *>* 10*µ*m, were modelled with a Generalized Additive Model (GAM, solid lines: living cells and dashed lines: dead and dying cells.). GAM is missing for dead and dying cells for 100*µ*E and 300*µ*E for diatoms with diameter *>* 10*µ*m due to too sparse data.

### Sinking patterns and departures from Stokes’ Law

Comparing the measured sinking velocities for growth days 1, 3, 5 and 7 for all combination of taxa, growth light condition, and viability with sinking velocities calculated according to Stokes’ Law (Equation 1 and 2 with information given in Table 2) showed large deviations from Stokes’ estimates (cf. Fig.4). In the case of studied cyanobacteria strains, the sinking velocity was greatly underestimated according to the Stokes’ approach independent of the growth light (cf. Fig.4), while in case of the studied diatoms strains the Stokes’ approach both under- and overestimated the sinking velocity. For growth light 300*µ*E only some measurements of *Thalassiosira weissflogii* and *Thalassiosira pseudonana* were in the same order as the Stokes’ estimate, while sinking velocities of *Minidiscus variabilis* were underestimated (cf. Fig.4C, F). For growth light 100*µ*E and 30*µ*E some measurements of *Thalassiosira pseudonana* and *Minidiscus variabilis* were in the same order as the Stokes’ estimate, while sinking velocities of *Thalassiosira weissflogii* were overestimated by the Stokes’ approach (cf. Fig.4A, B, D and E).

**Fig 4.**
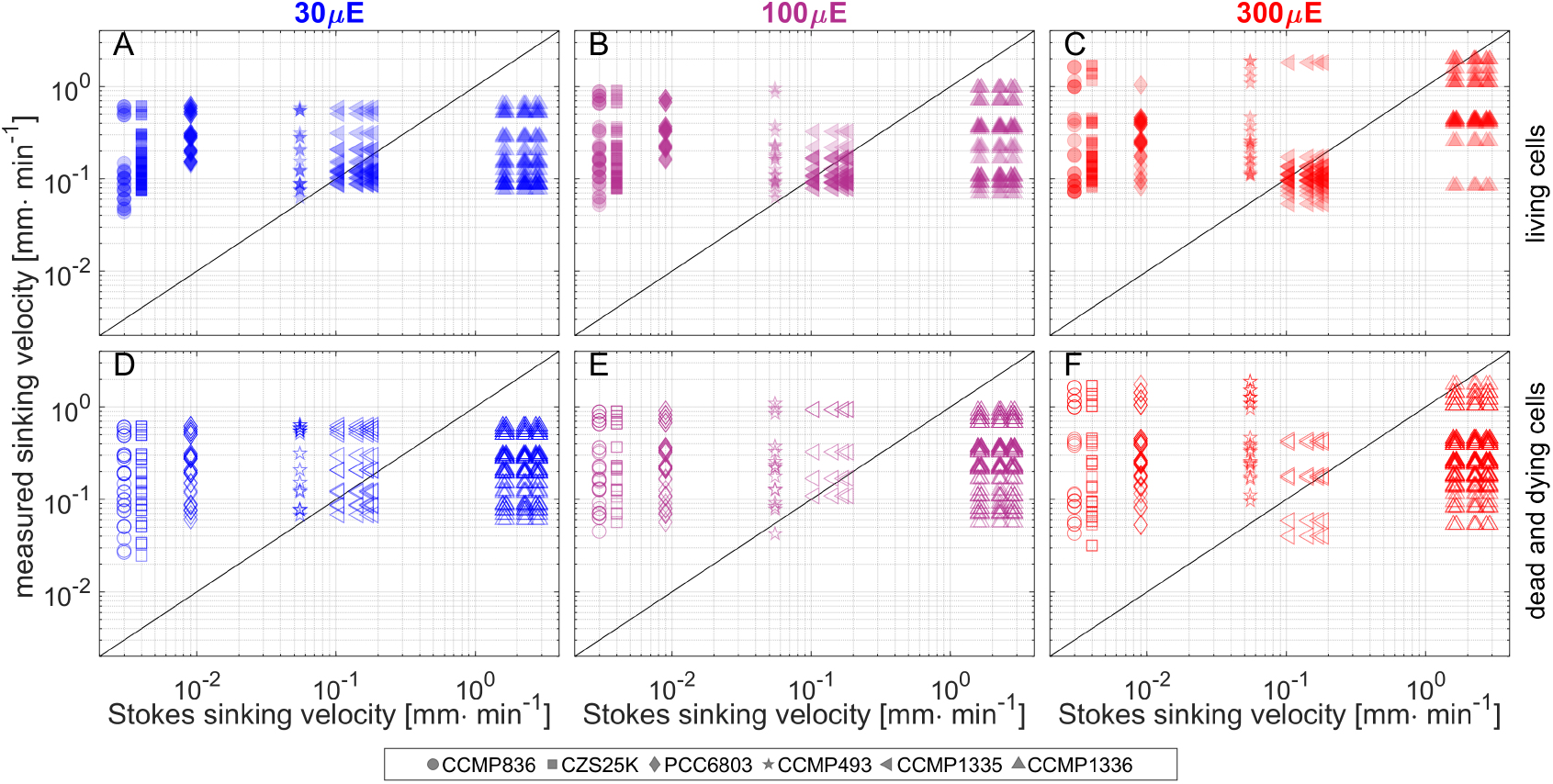
Relation of measured sinking velocity (pooled data for growth days 1, 3, 5 and 7) and sinking velocity calculated according to Stokes (Equation 1 and 2 with information given in Table 2) for living cells (A-C) and dead and dying cells (D-F) and growth light 30*µ*E (A, D), 100*µ*E (B, E) and 300*µ*E (C, F). For the cylindrical *Thalassiosira weissflogii* (CCMP1336) and *Thalassiosira pseudonana* (CCMP1335) several Stokes’ estimates are provided as different size information could be found in literature and both vertical as well as horizontal orientated sinking is possible. Filled symbols correspond to measurements of living cells and unfilled symbols to measurements of dead and dying cells. The black diagonals indicate the hypothetical case that Stokes sinking velocity and measured sinking velocity are equal. Cyanobacteria and diatom strains are abbreviated in the legend according to Table 1.

To approximate the potential impacts of different sinking velocities in an ocean setting we used data from the Baltic Sea as a test bench. Based on output from the ROMS-Bio-Optic model [48] and personal communication (B.Cahill), light levels of 300*µ*E, 100*µ*E and 30*µ*E in the western Baltic light attenuation field correspond to depths of about -2m, -5m, and -8m, respectively, during phytoplankton blooms.

We estimated the positional changes of cyanobacteria and diatoms over 24 hours, based on their respective measured sinking velocities, after starting at different depths equivalent to their growth light.

This approach focused on export in a simplified, turbulence-free environment to describe the potential impact of ‘only’ sinking within a day. Additional up- or downwelling [49–51], turbulence [27, 52] or eddy mediated processes [53–55] could be modelled based on daily and regional characteristics. Nonetheless, up- and downwelling in the selected area and time of the year ranged from +1 to -1m within 24 hours, similar to the magnitude of estimated diel sinking.

For Fig. 5 we pooled the results into cyanobacteria with diameter*<* 2.5*µ*m, diatoms with diameter *<* 10*µ*m, and diatoms with diameter *>* 10*µ*m based on the similar sinking properties within these groups. Both diatoms with diameter*<* 10*µ*m and cyanobacterial strains showed a split into fast and slow-sinking sub-populations, with differences in sinking displacement of up to 2m per day. Interestingly, the gap in diel displacement between fast and slow-sinking sub-populations increased with increasing light availability.

**Fig 5.**
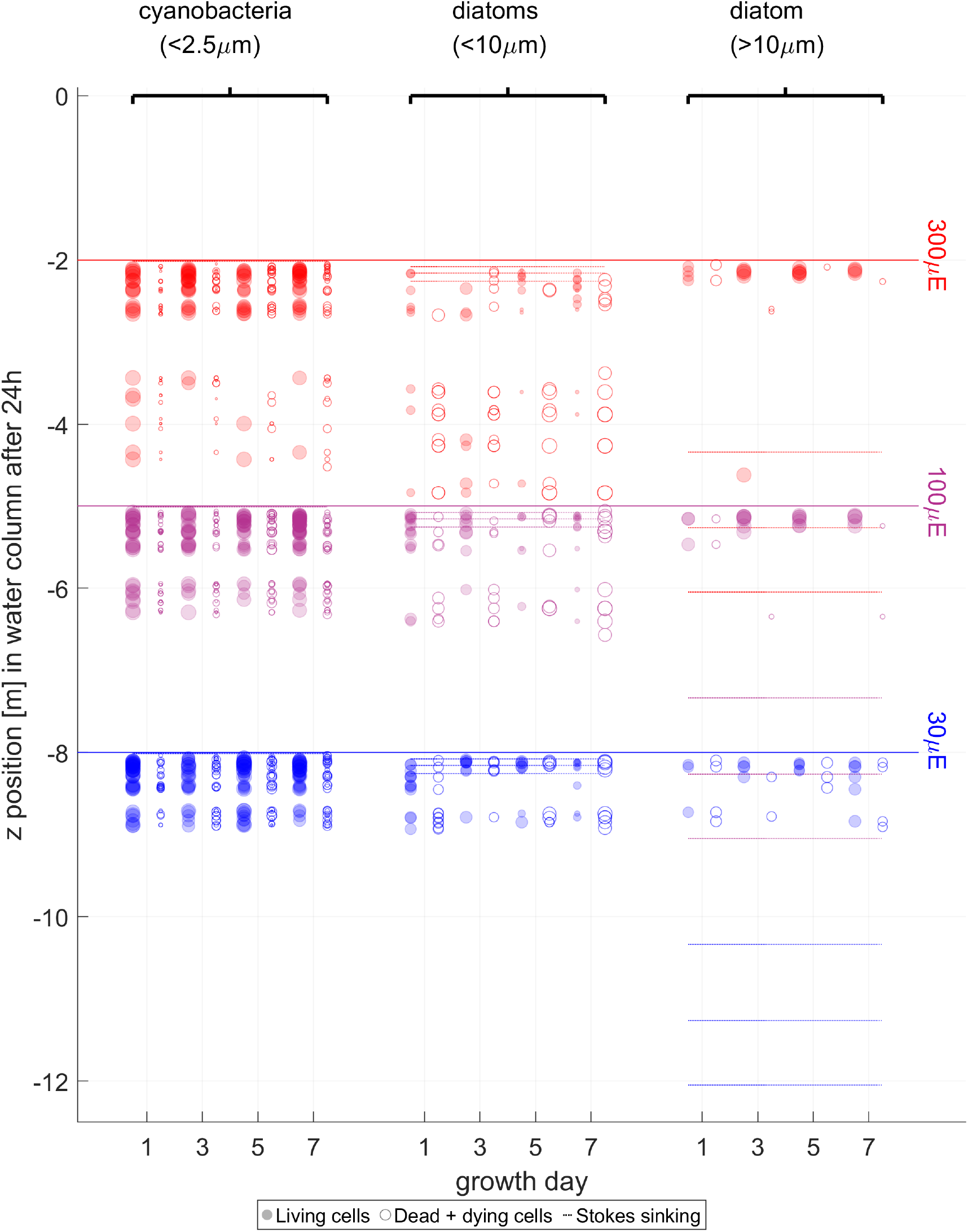
Cell positions after 24 hours sinking down a hypothetical water column, with single measurements of sinking velocities for cyanobacterial strains (*Synechococcus* sp., *Cyanobium* sp., *Synechocystis* sp.), diatoms *<* 10*µ*m (*Minidiscus variabilis, Thalassiosira pseudonana*) and a diatom *>* 10*µ*m (*Thalassiosira weissflogii* ) taken for their growth trajectory stage and viability state. The numbers in the brackets indicate the cell diameter. Cells start at different hypothetical depths, corresponding to their growth light level; the red solid line at 2m for 300*µ*E; purple line at 5m for 100*µ*E, and blue line at 8m for 30*µ*E. The filled circles correspond to living cells, the empty circles to the sum of dead and dying cells. Circles can overlap as part of several detected peaks corresponding to several sinking velocities per strain and condition, leading to darker filled circles. The circle size indicates the relative share per viability state of the total amount of cells from each strain and condition. The dotted horizontal lines show the sinking depths expected from a Stokes’ law calculation based upon cell radii (Eq. 1 and 2 and Table 2) for the different strains. The colors of the dotted lines correspond to the growth light climates, although the Stokes estimates do not depend directly upon growth light.

Additionally, Fig. 5 shows the relative share of living vs. dead and dying cells to the total count of cells at each point of the growth trajectory, as circle size.

In the cyanobacterial strains, living cyanobacterial cells dominated net-export at 300*µ*E early in the growth trajectory (Fig. 5). The share of dead and dying cyanobacteria cells, and thus their impact on net-export, increased over the growth trajectory, sinking up to 3m under 300*µ* from day 5 onwards. The trend of an increasing share of dead and dying cyanobacteria cells to net-export across the growth trajectory was also found at 100*µ*E and 30*µ*E, representative of deeper starting depths, but overall sinking displacement was much less under 30*µ*E.

In case of diatoms with diameter *<* 10*µ*m, the share of fast-sinking dead and dying cells was larger compared to living cells from the beginning of the growth trajectory and these dead and dying diatom cells often sank deeper than living diatoms cells within a day. Thus, living diatom cells with diameter *<* 10*µ*m would contribute less to net-export into deeper areas across all growth light levels, indicating either a change in sinking regulation or possibly a shift towards single cells rather than colonies. The share of fast-sinking, living diatoms with diameter *<* 10*µ*m was small compared to the total cells exported under 300*µ*E and 100*µ*E and was probably caused by chains or microcolonies of diatoms, rather than single cells.

In living diatoms with diameter *>* 10*µ*m no clear separation into fast- and slow-sinking sub-populations under higher growth light level (100 and 300*µ*E) could be observed, and overall vertical displacements were, surprisingly, smaller compared to the smaller phytoplankton groups in this study, and seemed independent of growth light.

A decomposition of predicted sinking displacements for strain can be found in Fig. S3.

For comparison we calculated sinking displacements within 24 hours using a simplified Stokes’ law [17, 22] (Equation 1 and 2 with information given in Table 2). As indicated by Fig.4 the depths reached according to Stokes’ (horizontal dotted lines in Fig.5) differed widely from our observed ones, with both over- and under-estimates of sinking depending upon the cell type.

Sinking distances over 24 hours estimated according to a simplified Stokes’ law for the cyanobacteria strains were negligible, in contrast to our estimated sinking distances of 1-2m over 24 hours, based upon measured sinking velocities of the cyanobacteria. Diatoms with diameter *<*10 *µ*m also showed a small sinking distance of only approximately 0.3m per day estimated according to Stokes, whereas, dead and dying diatoms cells *<*10 *µ*m actually sank much faster and deeper in our study, with estimated sinking sufficient that cells starting at depths equivalent to 300*µ*E could be carried all the way down to depths equivalent to 100*µ*E within a day (cf. Fig. 5). In marked contrast, for cells of the largest diatom (diameter *>* 10*µ*m) Stokes’ estimates of sinking velocities generated sinking of more than 2m per day, sufficient to move from 300*µ*E equivalent depth to 100*µ*E equivalent depth, 100*µ*E to 30*µ*E and from 30*µ*E to an aphotic depth within a day. But, using our measures of sinking velocities, neither living nor dead and dying diatom cells with diameter *>* 10*µ*m would actually sink that far. Stokes’ law based only on the cell size and assumptions of the cell mass density, even taking diatom silica walls into account, thus did not adequately describe sinking of monoclonal phytoplankton populations, heterogeneous in size and viability states in an oceanic water column, as pointed out by [29]. Differences in sinking velocities were influenced both by initial growth light, by cell viability state, taxa and somewhat by size class.

## Discussion

### Impact of growth stage and light on sinking

Our measured sinking velocities - with values between 0.1 and 1mm*·*min^−1^ for cyanobacteria and 0.1 and 1.8mm*·*min^−1^ for diatoms depending on size, viability state and light conditions (cf. Fig.2) - are in the same orders of magnitude reported in other studies e.g. [13, 14, 17, 24] and references in Table 2 in [46].

Sub-populations with different sinking velocities, within a monoclonal culture, could result from a combination of viability, intra-strain cell size differences, changes of cell shape and size during rapid growth as seen in *Synechocystis* sp. [56]. Additionally, the formation of cell colonies, as in case of cyanobacterial strains showing a trend towards cell colony formation along the growth trajectory, possibly as a stress reaction to high light, or through excretion of excess photosynthate ( [57] and reference cited therein), could explain the increase of sinking velocity for the fast-sinking sub-population of cyanobacteria along their growth trajectory (cf. Fig.3) under high growth light (300*µ*m). Under lower growth light (100*µ*m and 30*µ*m) this trend towards faster-sinking cells due to colony formation seems to be less pronounced (cf. Fig.3).

Our results likely reflect differing cell metabolic activities and compositions under different light climates, as hypothesized to influence sinking velocity by [33], who also discussed different cell shapes depending on growth light conditions, which would also affect sinking properties.

In the case of the diatoms *<*10 *µ*m increasing sinking velocities with increasing growth light may result from higher carbohydrate production under higher light, which increases the mass density of living, as well as dead cells. Similar changes of sinking velocity with growth stage were described for diatoms in an ocean setting [15, 33, 34].

These effects could also explain fluctuations in the sinking velocity for the fast-sinking sub-population along the growth trajectory of the diatoms with diameter *<* 10*µ*m: along the growth trajectory the cell size in the population increases, leading to a better ability to regulate sinking by metabolic processes. This growth-stage fluctuation is more pronounced under low light (cf. Fig.3).

In conclusion the convolution of age of the culture, growth light and viability produce a complex effects on sinking velocity dynamics that, differ for cyanobacteria and diatoms.

### Impacts of sinking on export

The different patterns of sinking extrapolated over 24 hours show the importance of viability-specific sinking velocities, as well as light climate and taxa-specific sinking, when estimating export of organic matter in the water column. Cells grown under high light conditions can increase organic matter content export to deeper layers, but will then sink more slowly upon reaching deeper areas as seen for cells with diameter *<* 10*µ*m. In contrast, cells grown under low light conditions tend to conserve the organic matter within the source light level depth, with their slower sinking. Taking into account hydrodynamical effects as upwelling and downwelling, the size abundance depth spectrum could change [49], influencing the biological carbon pump [2, 31].

A striking pattern was the increase of overall sinking velocity with increasing growth light level for small cells. This result implies that cells grown under low light are more buoyant and slow down organic matter transport to yet deeper layers. In our simplified export model we found that for export into deeper layers living cyanobacteria provided a larger share while for small diatoms (diameter *<* 10*µ*m) dead and dying cells play a more important role. As a result, fast-sinking living cyanobacteria growing under high light would export living biomass to deeper photic layers, while fast-sinking, dead and dying, small diatoms, growing under high light would promote remineralization processes in deeper layers. These results are consistent with [2], that at high latitudes in a community dominated by diatoms, only a small fraction of remineralization occurs in the surface layer.

Living large cells like *Thalassiosira weissflogii*, surprisingly, did not contribute to export into deeper layers but rather conserved their biomass in their photic layer of origin, probably through active regulation, since their dead and dying cells generally sank more.

Assuming an increasing fraction of primary production by small cells in surface layers due to climate change induced temperature rise (cf. [7, 9]), the upper layers will become more turbid. A consequence would be an increase in light attenuation in the upper layers, which probably leads to slower sinking cells, less likely to reach deeper layers and contribute to organic matter export, whether as living cyanobacteria, as dead small diatoms, or as sources for marine snow formation. This shift is supported by less wind-induced mixing and increased stratification in the ocean, which ultimately moves the layer of production up within the water column, possibly lowering carbon export into deeper waters. For deeper layers, more light attenuation above would mean slower delivery of organic matter content from shallower depths, due to the slowed sinking velocity. A potential implementation of our findings into ocean models would be the use of a light intensity dependent threshold-based switch between sinking velocities, as implemented in some freshwater ecosystem models [58].

In the global view, changing sinking velocities for cells traversing different light climates and growth stages could considerably affect our estimates of global carbon export.

In summary, unialgal cell suspensions show a wide range of sinking velocities across different sub-populations, which shift over growth trajectories. Sinking velocities also vary with growth light, with implications for export from the upper to deeper photic zone. Finally, the taxa show wide departures from Stoke’s law predictions of sinking, with implications for modelling.

## Supporting information

### Code and Data availability statement

The raw data will be deposited on Dryad upon acceptance. The code used to calculate the sinking velocities in this work is available openly and open source at GitHub https://github.com/maxberthold/PhytoplanktonSinkVelocities.

### Supporting information File

Additional information and figures can be found in Supporting information file.

## Acknowledgment

The authors would like to thank Bronwyn Cahill for providing information on the depths of light attenuation in the western Baltic Sea.

## Authors contribution

Conceptualization: DAC, MB, RV-K. Methodology: MB, RV-K. Investigation: MB, RV-K. Formal Analysis: MB, RV-K. Software: MB, RV-K. Visualization: MB, RV-K. Writing-original draft: MB, RV-K. Writing-review&editing: DAC, MB, RV-K. Resources: DAC.

## References

1. Pearre S. Eat and run? The hunger/satiation hypothesis in vertical migration: history, evidence and consequences. Biological Reviews of the Cambridge Philosophical Society. 2003;78(1):1–79. doi:10.1017/s146479310200595x.

2. Henson SA, Sanders R, Madsen E. Global patterns in efficiency of particulate organic carbon export and transfer to the deep ocean. Global Biogeochemical Cycles. 2012;26(1). doi:10.1029/2011gb004099.

3. Sabine CL, Feely RA, Gruber N, Key RM, Lee K, Bullister JL, et al. The Oceanic Sink for Anthropogenic CO2. Science. 2004;305(5682):367–371. doi:10.1126/science.1097403.

4. Pierce DW. Future Changes in Biological Activity in the North Pacific Due to Anthropogenic Forcing of the Physical Environment. Climatic Change. 2004;62(1-3):389–418. doi:10.1023/b:clim.0000013678.59224.98.

5. Sarmiento JL, Slater R, Barber R, Bopp L, Doney SC, Hirst AC, et al. Response of ocean ecosystems to climate warming. Global Biogeochemical Cycles. 2004;18(3):/a–n/a. doi:10.1029/2003gb002134.

6. Sommer U, Peter KH, Genitsaris S, Moustaka-Gouni M. Do marine phytoplankton follow Bergmann’s rule sensu lato? Biological Reviews. 2016;92(2):1011–1026. doi:10.1111/brv.12266.

7. Henson SA, Cael BB, Allen SR, Dutkiewicz S. Future phytoplankton diversity in a changing climate. Nature Communications. 2021;12(1). doi:10.1038/s41467-021-25699-w.

8. Richardson TL, Jackson GA. Small Phytoplankton and Carbon Export from the Surface Ocean. Science. 2007;315(5813):838–840. doi:10.1126/science.1133471.

9. Henson SA, Laufkötter C, Leung S, Giering SLC, Palevsky HI, Cavan EL. Uncertain response of ocean biological carbon export in a changing world. Nature Geoscience. 2022;15(4):248–254. doi:10.1038/s41561-022-00927-0.

10. Gemmell BJ, Oh G, Buskey EJ, Villareal TA. Dynamic sinking behaviour in marine phytoplankton: rapid changes in buoyancy may aid in nutrient uptake. Proceedings of the Royal Society B: Biological Sciences. 2016;283(1840):20161126. doi:10.1098/rspb.2016.1126.

11. Pfeifer F. Distribution, formation and regulation of gas vesicles. Nature Reviews Microbiology. 2012;10(10):705–715. doi:10.1038/nrmicro2834.

12. Schneegurt MA, Sherman DM, Nayar S, Sherman LA. Oscillating behavior of carbohydrate granule formation and dinitrogen fixation in the cyanobacterium Cyanothece sp. strain ATCC 51142. Journal of Bacteriology. 1994;176(6):1586–1597. doi:10.1128/jb.176.6.1586-1597.1994.

13. Bienfang PK. Sinking rates of heterogeneous, temperate phytoplankton populations. Journal of Plankton Research. 1981;3(2):235–253. doi:10.1093/plankt/3.2.235.

14. Strickland JDH. Measuring the production of marine phytoplankton. Fish Res Bd Canada Bull. 1960;122:172.

15. Smetacek VS. Role of sinking in diatom life-history cycles: ecological, evolutionary and geological significance. Marine Biology. 1985;84(3):239–251. doi:10.1007/bf00392493.

16. Walker M, Hammel JU, Wilde F, Hoehfurtner T, Humphries S, Schuech R. Estimation of sinking velocities using free-falling dynamically scaled models: foraminifera as a test case. Journal of Experimental Biology. 2020;doi:10.1242/jeb.230961.

17. Miklasz KA, Denny MW. Diatom sinkings speeds: Improved predictions and insight from a modified Stokes’ law. Limnology and Oceanography. 2010;55(6):2513–2525. doi:10.4319/lo.2010.55.6.2513.

18. Bannon CC, Campbell DA. Sinking towards destiny: High throughput measurement of phytoplankton sinking rates through time-resolved fluorescence plate spectroscopy. PLOS ONE. 2017;12(10):e0185166. doi:10.1371/journal.pone.0185166.

19. Bienfang PK, Harrison PJ, Quarmby LM. Sinking rate response to depletion of nitrate, phosphate and silicate in four marine diatoms. Marine Biology. 1982;67(3):295–302. doi:10.1007/bf00397670.

20. Wang X, Sun J, Wei Y, Wu X. Response of the Phytoplankton Sinking Rate to Community Structure and Environmental Factors in the Eastern Indian Ocean. Plants. 2022;11(12):1534. doi:10.3390/plants11121534.

21. Bach LT, Riebesell U, Sett S, Febiri S, Rzepka P, Schulz KG. An approach for particle sinking velocity measurements in the 3-400μm size range and considerations on the effect of temperature on sinking rates. Marine Biology. 2012;159(8):1853–1864. doi:10.1007/s00227-012-1945-2.

22. Walsby AE, Holland DP. Sinking velocities of phytoplankton measured on a stable density gradient by laser scanning. Journal of The Royal Society Interface. 2005;3(8):429–439. doi:10.1098/rsif.2005.0106.

23. Shimoda Y, Arhonditsis GB. Phytoplankton functional type modelling: Running before we can walk? A critical evaluation of the current state of knowledge. Ecological Modelling. 2016;320:29–43. doi:10.1016/j.ecolmodel.2015.08.029.

24. Riley GA. Quantitative ecology of the plankton of the western North Atlantic. Bull Bingham Oceanogr Collection. 1949;12:1–169.

25. Stemmann L, Jackson GA, Ianson D. A vertical model of particle size distributions and fluxes in the midwater column that includes biological and physical processes—Part I: model formulation. Deep Sea Research Part I: Oceanographic Research Papers. 2004;51(7):865–884. doi:10.1016/j.dsr.2004.03.001.

26. Stemmann L, Jackson GA, Gorsky G. A vertical model of particle size distributions and fluxes in the midwater column that includes biological and physical processes—Part II: application to a three year survey in the NW Mediterranean Sea. Deep Sea Research Part I: Oceanographic Research Papers. 2004;51(7):885–908. doi:10.1016/j.dsr.2004.03.002.

27. Ruiz J, Macías D, Peters F. Turbulence increases the average settling velocity of phytoplankton cells. Proceedings of the National Academy of Sciences. 2004;101(51):17720–17724. doi:10.1073/pnas.0401539101.

28. Monroy P, Hernández-García E, Rossi V, López C. Modeling the dynamical sinking of biogenic particles in oceanic flow. Nonlinear Processes in Geophysics. 2017;24(2):293–305. doi:10.5194/npg-24-293-2017.

29. McDonnell AMP, Buesseler KO. Variability in the average sinking velocity of marine particles. Limnology and Oceanography. 2010;55(5):2085–2096. doi:10.4319/lo.2010.55.5.2085.

30. Bissett WP, Walsh JJ, Dieterle DA, Carder KL. Carbon cycling in the upper waters of the Sargasso Sea: I. Numerical simulation of differential carbon and nitrogen fluxes. Deep Sea Research Part I: Oceanographic Research Papers. 1999;46(2):205–269. doi:10.1016/s0967-0637(98)00062-4.

31. Manizza M, Quéré CL, Watson AJ, Buitenhuis ET. Ocean biogeochemical response to phytoplankton-light feedback in a global model. Journal of Geophysical Research. 2008;113(C10). doi:10.1029/2007jc004478.

32. Neumann T, Radtke H, Cahill B, Schmidt M, Rehder G. Non-Redfieldian carbon model for the Baltic Sea (ERGOM version 1.2) - implementation and budget estimates. Geoscientific Model Development. 2022;15(22):8473–8540. doi:10.5194/gmd-15-8473-2022.

33. Riley GA. Physiological aspects of spring diatom flowerings. Peabody Mus. of Natural History, Yale Univ.; 1943.

34. Smayda TJ. Normal and accelerated sinking of phytoplankton in the sea. Marine Geology. 1971;11(2):105–122. doi:10.1016/0025-3227(71)90070-3.

35. Field CB, Behrenfeld MJ, Randerson JT, Falkowski P. Primary Production of the Biosphere: Integrating Terrestrial and Oceanic Components. Science. 1998;281(5374):237–240. doi:10.1126/science.281.5374.237.

36. Flombaum P, Gallegos JL, Gordillo RA, Rincón J, Zabala LL, Jiao N, et al. Present and future global distributions of the marine Cyanobacteria Prochlorococcus and Synechococcus. Proceedings of the National Academy of Sciences. 2013;110(24):9824–9829. doi:10.1073/pnas.1307701110.

37. Zeileis A, Leisch F, Hornik K, Kleiber C. strucchange: An R Package for Testing for Structural Change in Linear Regression Models. Journal of Statistical Software. 2002;7(2):1–38. doi:10.18637/jss.v007.i02.

38. Williams SC, Hong Y, Danavall DCA, Howard-Jones MH, Gibson D, Frischer ME, et al. Distinguishing between living and nonliving bacteria: Evaluation of the vital stain propidium iodide and its combined use with molecular probes in aquatic samples. Journal of Microbiological Methods. 1998;32(3):225–236. doi:10.1016/s0167-7012(98)00014-1.

39. R Core Team. R: A Language and Environment for Statistical Computing; 2021. Available from: https://www.R-project.org/.

40. Hildebrand M, York E, Kelz JI, Davis AK, Frigeri LG, Allison DP, et al. Nanoscale control of silica morphology and three-dimensional structure during diatom cell wall formation. Journal of Materials Research. 2006;21(10):2689–2698. doi:10.1557/jmr.2006.0333.

41. Olenina I, Hajdu S, Edler L, Andersson A, Wasmund N, Busch S, et al. Biovolumes and size-classes of phytoplankton in the Baltic Sea. HELCOM Balt Sea Environ Proc No 106, 144 pp. 2006;.

42. Hildebrand M, Holton G, Joy DC, Doktycz MJ, Allison DP. Diverse and conserved nano- and mesoscale structures of diatom silica revealed by atomic force microscopy. Journal of Microscopy. 2009;235(2):172–187. doi:10.1111/j.1365-2818.2009.03198.x.

43. Kaczmarska I, Lovejoy C, Potvin M, Macgillivary M. Morphological and molecular characteristics of selected species of Minidiscus (Bacillariophyta, Thalassiosiraceae). European Journal of Phycology. 2009;44(4):461–475. doi:10.1080/09670260902855873.

44. Leppäranta M, Myrberg K. Physical oceanography of the Baltic Sea. Praxis Publishing. Springer: Berlin, Heidelberg, New York; 2009.

45. Viswanath DS, Natavajan G. Data book on the viscosity of liquids. New York, NY (USA); Hemisphere Publishing. 1989;.

46. Qiu Z, Doglioli AM, Carlotti F. Using a Lagrangian model to estimate source regions of particles in sediment traps. Science China Earth Sciences. 2014;57(10):2447–2456. doi:10.1007/s11430-014-4880-x.

47. Arrieta J, Jeanneret R, Roig P, Tuval I. On the fate of sinking diatoms: the transport of active buoyancy-regulating cells in the ocean. Philosophical Transactions of the Royal Society A: Mathematical, Physical and Engineering Sciences. 2020;378(2179):20190529. doi:10.1098/rsta.2019.0529.

48. Cahill BE, Kowalczuk P, Kritten L, Gräwe U, Wilkin J, Fischer J. Estimating the seasonal impact of optically significant water constituents on surface heating rates in the western Baltic Sea. Biogeosciences. 2023;20(13):2743–2768. doi:10.5194/bg-20-2743-2023.

49. Rodríguez J, Tintoré J, Allen JT, Blanco JM, Gomis D, Reul A, et al. Mesoscale vertical motion and the size structure of phytoplankton in the ocean. Nature. 2001;410(6826):360–363. doi:10.1038/35066560.

50. Rossi V, López C, Sudre J, Hernández-García E, Garçon V. Comparative study of mixing and biological activity of the Benguela and Canary upwelling systems. Geophysical Research Letters. 2008;35(11). doi:10.1029/2008gl033610.

51. Gruber N, Lachkar Z, Frenzel H, Marchesiello P, Münnich M, McWilliams JC, et al. Eddy-induced reduction of biological production in eastern boundary upwelling systems. Nature Geoscience. 2011;4(11):787–792. doi:10.1038/ngeo1273.

52. Bracco A, Provenzale A, Scheuring I. Mesoscale vortices and the paradox of the plankton. Proceedings of the Royal Society of London Series B: Biological Sciences. 2000;267(1454):1795–1800. doi:10.1098/rspb.2000.1212.

53. McGillicuddy DJ. Mechanisms of Physical-Biological-Biogeochemical Interaction at the Oceanic Mesoscale. Annual Review of Marine Science. 2016;8(1):125–159. doi:10.1146/annurev-marine-010814-015606.

54. Vortmeyer-Kley R, Lünsmann B, Berthold M, Gräwe U, Feudel U. Eddies: Fluid Dynamical Niches or Transporters?–A Case Study in the Western Baltic Sea. Frontiers in Marine Science. 2019;6. doi:10.3389/fmars.2019.00118.

55. Freilich MA, Flierl G, Mahadevan A. Diversity of Growth Rates Maximizes Phytoplankton Productivity in an Eddying Ocean. Geophysical Research Letters. 2022;49(3). doi:10.1029/2021gl096180.

56. Schuurmans RM, Matthijs JCP, Hellingwerf KJ. Transition from exponential to linear photoautotrophic growth changes the physiology of Synechocystis sp. PCC 6803. Photosynthesis Research. 2017;132(1):69–82. doi:10.1007/s11120-016-0329-8.

57. Callieri C, Cronberg G, Stockner JG. Freshwater Picocyanobacteria: Single Cells, Microcolonies and Colonial Forms. In: Ecology of Cyanobacteria II. Springer Netherlands; 2012. p. 229–269.

58. Reynolds CS. The ecology of phytoplankton. Cambridge University Press; 2006.

